# PKN2 regulates cell-junctions to limit colitis

**DOI:** 10.64898/2025.12.15.694339

**Authors:** Jack C Henry, Ashley Campbell, Prerana Huddar, Joanne Sewell, Shinelle Menezes, Adam Passman, Kane Smith, Shannah McGauran, Ivan Quétier, Meng-Lay Lin, Marnix Jansen, Neil McDonald, Trevor A Graham, Stuart McDonald, Angus JM Cameron

## Abstract

Intestinal barrier failure is a defining feature of inflammatory bowel disease and a key driver of inflammation-associated colorectal cancer. However, the epithelial mechanisms that preserve barrier stability during inflammatory stress remain incompletely defined. Here, we identify the Rho-regulated kinase PKN2 as a non-redundant safeguard of tight junction integrity and a candidate tumour suppressor in colitis-associated colorectal cancer. Conditional PKN2 deletion in mice sensitised the colon to inflammatory injury and significantly increased adenoma formation in an inflammatory cancer model, with tumour burden tightly correlating with colitis severity. Mechanistically, PKN2 localises to tight junctions and is required to stabilise barrier integrity during injury in mouse and organoid models. PKN2 loss induces transcriptional programs conserved in human inflammatory bowel disease, where reduced PKN2 expression associates with increased disease severity and altered therapeutic response. Notably, PKN2 is encoded within genomic loci previously linked to intestinal inflammation susceptibility and tumour suppression in both humans and mice. Our findings identify PKN2 as a functional effector underlying these phenotypes and demonstrate that even heterozygous loss is sufficient to confer heightened sensitivity to inflammatory injury and tumour initiation, positioning PKN2 as a central regulator of tight junction stability that shapes disease severity, treatment response and cancer risk.

## Introduction

Inflammatory bowel disease (IBD), comprising Crohn’s disease and ulcerative colitis, is a group of chronic inflammatory disorders of the gastrointestinal tract, with a prevalence of approximately 0.3% in Western populations and a rising incidence in newly industrialized countries^1^. It is well established that patients with IBD have a significantly higher risk of developing colorectal cancer compared to the general population and therefore often undergo regular colorectal screening tests to detect and remove pre-malignant lesions^2-4^. Chronic inflammation plays a central role in the development and progression of many cancers, including both sporadic and colitis-associated colorectal cancer. Supporting this, anti-inflammatory drugs such as Aspirin and COX inhibitors have been shown to significantly reduce the risk of developing sporadic colorectal cancer^5-8^. In experimental models, disrupting inflammatory pathways, both intrinsic to epithelial cells and through immune cell targeting, reduces tumour formation in both the azoxymethane/dextran sodium sulphate (AOM/DSS) and genetic APC-targeting models^9-12^.

Given the strong link between inflammation and colorectal cancer risk, there has been considerable interest in identifying genetic loci that predispose individuals to IBD. Chromosome arm 1p has been identified as a significant susceptibility locus for human IBD; however, it contains over a thousand genes, making it difficult to pinpoint candidates^13^. Two potential candidates on 1p36 include TNF receptor and Caspase 9, yet other candidates on 1p are yet to be discovered^14,15^. Genetic mouse studies have implicated regions on chromosome 1p by exploring differential sensitivity in different background strains. *Cdcs1 (cytokine deficiency-induced colitis susceptibility-1)* on chromosome 3 is a locus containing 340 genes, syntenic with human 1p. Cdcs1 is well characterised as the cause of increased susceptibility between different mouse strains in several knockout colitis models^16^. Further research also identified the overlapping (colitis-associated) colon cancer susceptibility locus on mouse chromosome 3 (CCs3), which drives increased susceptibility to AOM/DSS-induced colorectal tumours in the A/J mouse strain when compared to C57BL/6J^17^.

Here, we identify PKN2 as a key colitis susceptibility gene on human chromosome 1p, syntenic with murine *Cdcs1* and CCs3. Global or epithelial-specific loss of PKN2 in vivo sensitises mice to DSS-induced colonic injury and promotes colorectal adenoma formation. Mechanistically, PKN2 deficiency disrupts epithelial tight junction integrity and barrier permeability, providing new insight into the pathogenesis of colitis and colorectal cancer.

## Results

### PKN2 loss promotes colitis and colitis-associated colorectal cancer

To test if PKN2 can act as a colitis-associated tumour suppressor, we exploited our inducible conditional PKN2 knockout mouse (global-iPKN2^KO^: *Pkn2*^*fl/fl*^*;Rosa26-CreER*^*T2*^) and the two-stage azoxymethane (AOM)/Dextran sodium-sulphate (DSS) colitis-associated colorectal tumour model (Fig. 1A). Global PKN2^KO^ is embryonic lethal^18^, but inducible conditional deletion in adults is generally well-tolerated^18,19^ with minimal macroscopic effects on normal colon homeostasis. Five consecutive days of treatment with tamoxifen in the iPKN2 model was sufficient to activate Cre and induce loss of PKN2 DNA and protein in the intestines of adult mice (Supplementary Fig. S1A, B) ^18^. To induce tumours, WT and PKN2 targeted mice were treated with a single dose of the mutagen AOM (10mg/kg) followed by three repeated rounds of 1% DSS, which typically results in tumour formation at 90 days^20^. Typically, DSS is administered at 2-2.5% in C57BL/6 mice but pilot experiments indicated that PKN2 targeted mice tolerated 2% DSS regimens poorly (Supplementary Fig. S1C-E). Even at the reduced dose of 1% DSS, both heterozygous (global-iPKN2^HET^) and homozygous PKN2 (global-iPKN2^KO^) deletion significantly exacerbated weight loss, disease activity and survival (scored 1-14) ^21^ (Fig. 1B-G). Surviving mice with PKN2 loss continued to show increased weight loss and disease activity across the experimental time course (Supplementary Fig. S1F-I). At 90-days post-AOM, mice were examined for tumour induction, which revealed an increase in macroscopically visible tumour incidence, multiplicity and tumour burden in PKN2-targeted mice (Fig. 1H-K). No tumours were observed at 90 days in cohorts treated with AOM alone. Pathological analysis identified that global-iPKN2^KO^ bowels presented with more high-grade tubular adenomas, while incidence of low-grade tissue dysplasia and tumour size distribution was similar between genotypes (Supplementary Fig. S1J-L). Globally targeting PKN2 did not induce polydipsia or alter relative DSS dose (Supplementary Fig. S1M). PKN2 targeted mice also presented with splenomegaly (Supplementary Fig. S1N, O) but had no change in colon length or overt pathology associated with other organs (Supplementary Fig. S1P). We conclude that haploinsufficiency of PKN2 is sufficient to exacerbate DSS induced disease and colorectal cancer adenoma formation, and that homozygous loss induces a more penetrant phenotype.

**Fig. 1.**
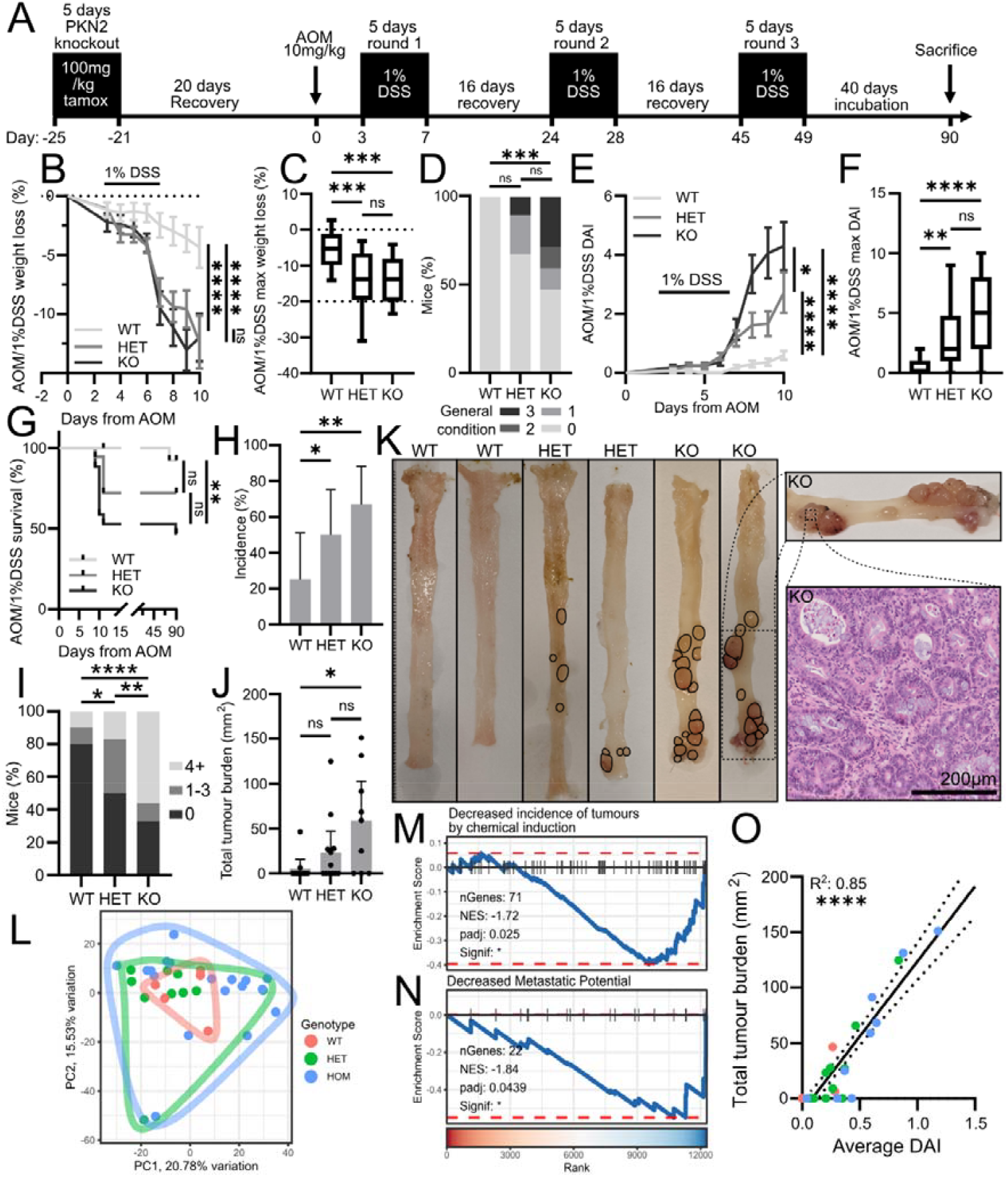
PKN2 loss sensitises mice to AOM/DSS treatment and promotes colitis-associated colorectal cancer. **(A)** Schematic of the AOM/1% DSS protocol. **(B, C)** Average (B) and maximum (C) weight change during the first 10 days of AOM/DSS treatment (WT: n=17, HET: n=18, KO: n=17; two-way RM-ANOVA (B), Tukey’s test (C). **(D)** General condition scores (0-3) during the first 10 days of treatment (WT: n=17, HET: n=18, KO: n=17; Dunnett’s test). **(E, F)** Average (E) and maximum (F) disease activity index (DAI) (WT: n=17, HET: n=18, KO: n=17; two-way RM-ANOVA (E), Tukey’s test (F)). **G)** Kaplan Meier survival analysis (WT: n=17, HET: n=18, KO: n=17; Log-rank test). *(H-J)* Adenoma incidence (H), multiplicity (J) and burden (K) at day 90 (WT: n=10, HET: n=12, KO: n=9; χ^2^ test (H, I); Dunn’s test (J)). **(K)** Representative colon images following the AOM1%DSS model with example pathology. **(L)** PCA of Smart3-SEQ epithelial samples (WT: n=5, HET: n=10, KO: n=18 adenomas). **(M, (N)** Gene set enrichment analysis of KO vs WT adenomas using mouse tumour phenotypes ontology gene sets. **(O)** Correlation of tumour burden and average disease activity (WT: n=10, HET: n=12, KO: n=9).

We sought to characterise the adenomas to identify phenotypes associated with PKN2 loss. Immunohistochemical analysis of immune cell markers and ki67 expression identified no obvious differences in immune cell infiltrate or proliferation between genotypes (Supplementary Fig. S2A-E). To further explore the adenoma phenotypes, we performed Smart-3SEQ transcriptomics^22^ on a selection of laser-captured epithelial regions for each genotype. Importantly, PKN2 expression is significantly reduced in heterozygous and homozygous knockouts of PKN2 (Supplementary Fig. S2F). However, principal component analysis identifies largely overlapping clustering between genotypes (Fig. 1L), reflecting a random, varied tumour population in which PKN2 status does not consistently dictate selected tumour phenotypes. In addition, gene set enrichment analysis between wild-type and global-iPKN2^KO^ adenomas does not identify any clear increase in cancer hallmarks or oncogenic signalling pathway activity, including APC/WNT, RAS, TP53 and YAP (Supplementary Fig. S2G, H). One of the most significant negatively enriched gene sets are genes upregulated by a constitutively active (Q63L) form of RHOA, a regulator of PKN2. Interestingly, six pathways were also significantly enriched in KO adenomas from the mouse tumour phenotypes ontology collection of gene sets, including tumour incidence by chemical induction, tumour growth size and metastatic potential, suggesting PKN2 might antagonise the expression of pro-tumorigenic genes in this model (Fig. 1M, N, S4I).

The increased sensitivity to DSS and high-grade dysplasia incidence, coupled with largely similar adenoma transcriptomic profiles, led us to hypothesise that PKN2 loss may act primarily by exacerbating a pro-neoplastic environment. This is further supported by exquisitely tight correlation between tumour burden and average disease activity (Fig. 1O).

### PKN2 loss aggravates inflammatory bowel injury

We next examined the impact of PKN2 loss on acute DSS treatment. Bowels harvested 8-days post initiation of 1% DSS treatment presented regions of epithelial erosion, exudate, goblet cell depletion and crypt damage with the most severe phenotypes associated with PKN2 loss (Fig. 2A-E).

**Fig. 2.**
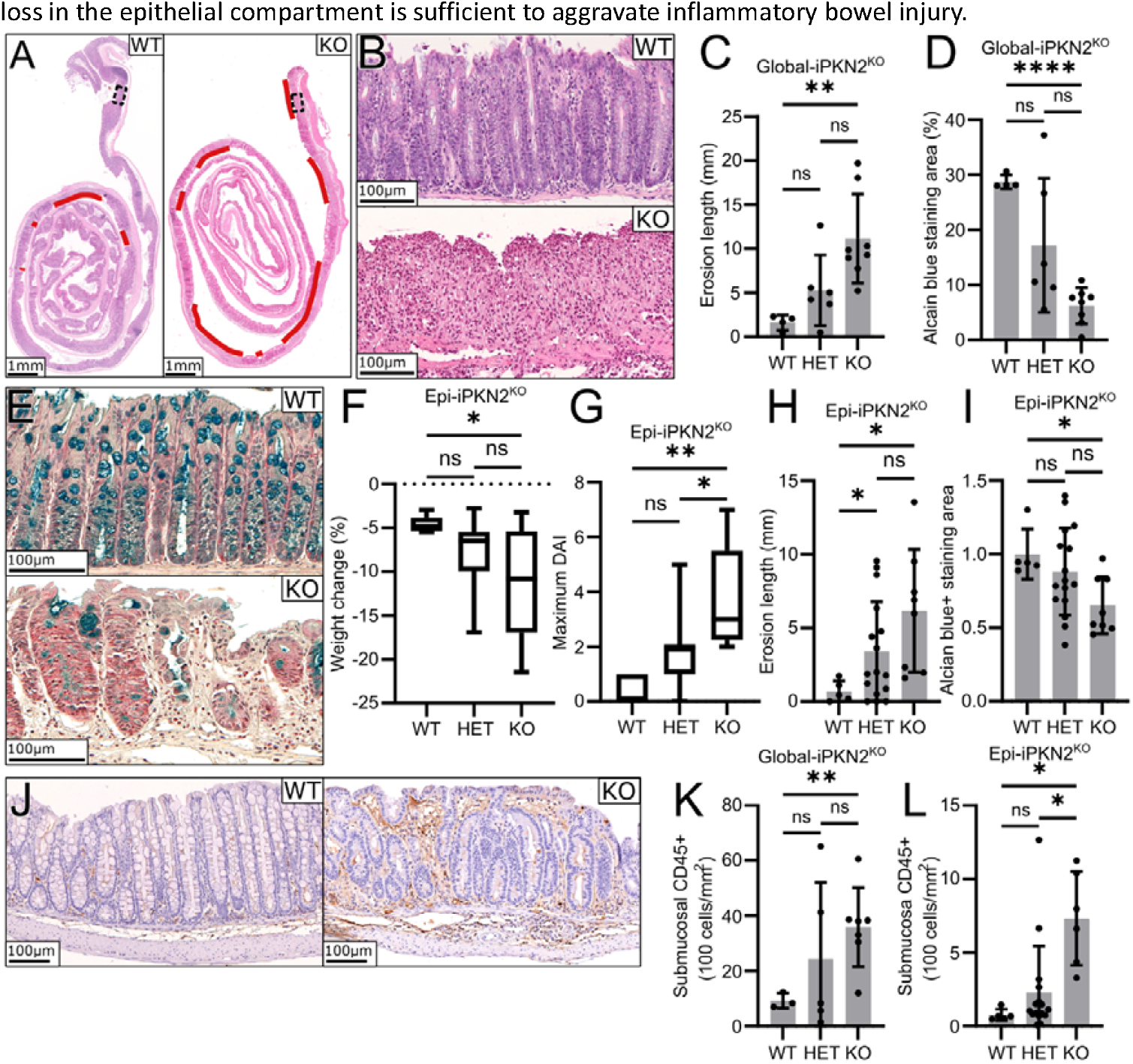
Epithelial PKN2 loss aggravates inflammatory bowel injury. **(A-C)** Epithelial erosion 8 days after AOM/1%DSS treatment in global-iPKN2^KO^ mice (WT: n=4, HET: n=6, KO: n=8; Dunnett’s test). Erosions are indicated in red and region shown in (B) is expanded in (A). **(D, E)** Goblet cell depletion assessed by alcian blue staining following AOM/1%DSS treatment in global-iPKN2^KO^ mice (WT: n=4, HET: n=6, KO: n=8; Dunnett’s test). **(F, G)** Maximum weight loss and disease activity index (DAI) during the first 10 days of 2%DSS treatment in epi-iPKN2^KO^ mice (WT: n=5, HET: n=16, KO: n=8; Tukey’s test). **(H, I)** Erosion length (H) and goblet cell depletion following 2%DSS treatment in epi-iPKN2^KO^ mice (WT: n=5, HET: n=16, KO: n=8; Dunnett’s test). **(J-L)** CD45+ immune cell infiltration into distal submucosa in global-iPKN2^KO^ mice following AOM/1% DSS treatment (J, K; WT: n=3, HET: n=5 KO: n=7) and epi-iPKN2^KO^ mice in response to 2%DSS treatment (L; WT: n=5, HET: n=16, KO: n=6; Dunnett’s test).

Single-cell RNA sequencing studies of normal and Crohn’s disease inflamed human colon identify PKN2 expression is enriched in, but not restricted to, epithelial cells (Supplementary Fig. S3A, B). To explore if sensitisation to inflammatory bowel injury reflects an epithelial-specific role for PKN2, the PKN2^flox^ mouse was crossed onto the Villin-Cre (*Pkn2*^*fl/fl*^*;Vil1-CreER*^*T2*^) specific driver, allowing for intestinal epithelial-specific knockout of PKN2 (Epi-iPKN2^KO^). PCR analysis of isolated epithelial crypts two weeks following tamoxifen treatment identified strong PKN2 locus recombination in the epithelial compartment of epi-iPKN2^KO^ mice (Supplementary Fig. S3C). Following a 2% DSS treatment regime without AOM, epi-PKN2^KO^ mice show a similar sensitisation to the global-PKN2^KO^, with exacerbated weight loss and DAI scores (Fig. 2F, G). Bowel tissue harvested 8-days post 2% DSS treatment also presented with a significant decrease in total colon length (Supplementary Fig. S3D) but no change to spleen size (Supplementary Fig. S3E). Histologically, mice with epithelial-specific PKN2 loss also had an increased prevalence of erosions, crypt loss and depletion of goblet cells (Fig. 2H, 2I, S3F, S3G). Erosions were primarily localised to distal regions in all genotypes. On average, distal submucosae showed increased thickness in global and epi-iPKN2^KO^ mice with higher levels of cellularity and CD45+ immune cell infiltrate (Fig. 2J-L, S3H-L). Surrounding distal crypts also contained more CD45+ immune cell infiltrate, indicating heightened local inflammation (Supplementary Fig. S3M, N). These analyses indicate that either homozygous or heterozygous PKN2 loss in the epithelial compartment is sufficient to aggravate inflammatory bowel injury.

### PKN2 loss perturbs ZO-1 tight junctions and models’ human colitis

To investigate epithelial phenotypes in vitro, colonic organoids were generated from wild-type C57BL/6 and *Pkn2*^*fl/fl*^*;Rosa26-CreER*^*T2*^ mice, enabling the inducible deletion of PKN2 with 4-hydroxytamoxifen (TAM) treatment (Supplementary Fig. S4A). Wild-type organoids treated with tamoxifen served as controls. RNA sequencing revealed that PKN2 loss was the dominant source of transcriptional variance, with tight clustering of biological replicates by principal component analysis (Supplementary Fig. S4B), PKN2 is the most significantly downregulated gene, confirming penetrant deletion (Supplementary Fig. S4C). While differential expression analysis identified modulation of several cancer-associated genes, PKN2-deficient organoids showed no evidence of spontaneous transformation (Supplementary Fig. S4D). Comparison with human IBD biopsies and colitis-derived organoids revealed substantial overlap in hallmark GSEA signatures, indicating that epithelial PKN2 loss recapitulates key transcriptional features of human colitis (Fig. 3A).

**Fig. 3.**
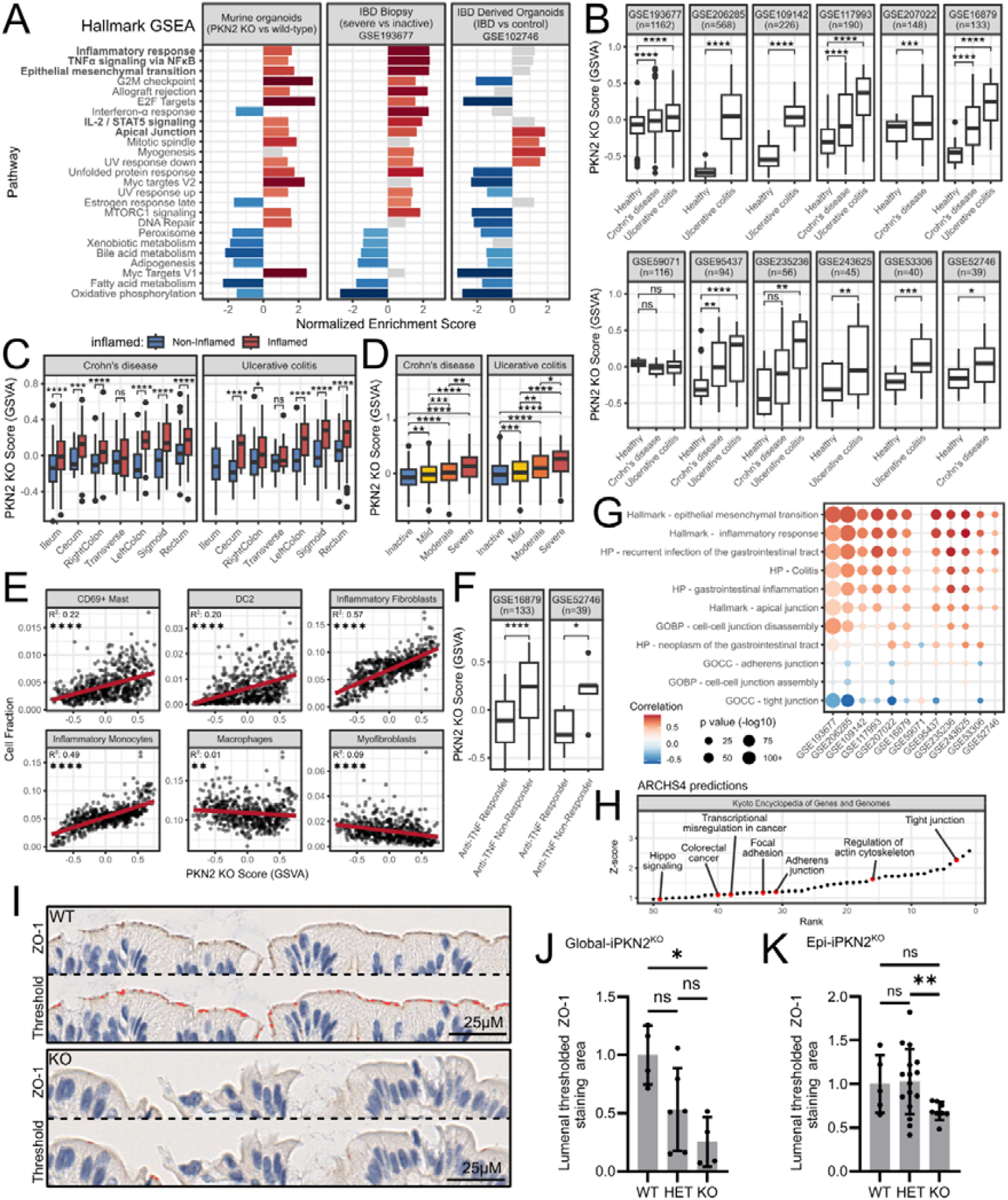
Transcriptomic profiling of PKN2 loss reveals inflammatory and junctional pathway modulation. **(A)** Hallmark gene set enrichment analysis comparing transcriptional responses to PKN2^KO^ in organoids with differential expression in patients with severe versus inactive IBD and primary colitis organoids versus healthy controls. **(B)** PKN2^KO^ score in patients with Crohn’s and ulcerative colitis compared with healthy controls across 12 independent bulk RNA-sequencing and microarray datasets. **(C, D)** PKN2^KO^ score in inflamed versus non-inflamed bowel regions (C) and across disease severity defined by endoscopic assessment (D). **(E)** Correlation between PKN2^KO^ score and immune cell infiltration; line of best fit, R^2^ and significance levels shown. **(F)** PKN2^KO^ score in Anti-TNF therapy responders and non-responders. **(G)** Correlation dot plot comparing PKN2^KO^ score with selected gene set enrichment by GSVA. **(H)** Top 50 ranked KEGG gene sets predicting PKN2 function identified by PrismEXP across the ARCHS4 compendium. **(I-K)** Immunohistochemical analysis of distal luminal epithelial ZO-1 staining in mice 8 days following 1% DSS treatment in global-iPKN2^KO^ (I, J; WT: n=4, HET: n=6, KO: n=4) and epithelial-iPKN2^KO^ mice (K; WT: n=4, HET: n=6, KO: n=4; Holm-Šídák’s test). Pixel threshold DAB staining intensity shown bottom.

Using our PKN2^KO^ epithelial transcriptional signatures, we applied PrismEXP predictions across the ARCHS4 compendium of integrated RNA-sequencing datasets to assess associations with human disease phenotypes. This revealed significant links to rectal prolapse, hematochezia and colon cancer (Supplementary Fig. S4E), all of which were exacerbated in our PKN2^KO^ mouse models. To relate these phenotype associations to human epithelial transcriptional changes, we derived a PKN2^KO^ epithelial signature from differentially expressed genes in PKN2-deficient organoids. Bulk PKN2 expression showed inverse correlation with this signature, consistent with reduced PKN2 expression driving a PKN2^KO^ transcriptional program (Supplementary Fig. S4F). Across 11 of 12 independent human IBD cohorts, the PKN2^KO^ signature was significantly elevated in both Ulcerative colitis and Crohn’s disease compared with healthy controls (Fig. 3B). Signature scores were higher in active versus inactive disease, enriched in inflamed compared with non-inflamed regions and maximal in distal colonic samples, mirroring observations in mouse models (Fig. 3C, S4G). Notably, PKN2^KO^ scores were highest in patients with severe disease, linking loss of epithelial *PKN2* activity to disease severity and highlighting its potential importance in driving clinically aggressive inflammatory pathology in human IBD (Fig. 3D).

Phenotypically, samples exhibiting an epithelial PKN2 loss-like signature showed expansion of inflammatory monocytes, fibroblasts, dendritic cells and mast cell fractions, indicating a shift toward an activated innate immune and stromal microenvironment (Fig. 3E). Minimal correlation with macrophage and myofibroblast populations was observed, suggesting a stronger association with active inflammation over tissue repair (Fig. 3E). Concordantly, GSEA revealed amplification of TNFα– NFκB and IL2-STAT5 driven inflammatory programs that sustain pathogenic T-cell responses and promote non-resolving mucosal injury (Fig. 3G). Notably, the PKN2^KO^ score also predicted response to anti-TNF therapy, linking epithelial PKN2 loss to clinically actionable inflammatory states (Fig. 3F). GSVA also demonstrated associations with inflammation, colitis and recurrent infection gene sets. Limited enrichment of gene sets involved in neoplasm-related programs further suggests that PKN2 loss promotes inflammatory damage-linked tumorigenesis rather than driving direct oncogenic programs, consistent with our in vivo findings (Fig. 3G).

To gain some mechanistic insight into these transcriptional and clinical associations, we applied PrismEXP analysis on KEGG gene sets and found that PKN2 expression is strongly linked to cell junction organisation, mechanotransduction and transcriptional misregulation (Fig. 3H). Consistent with this, hallmark apical junction gene sets were concordantly enriched in PKN2^KO^ organoids, IBD biopsies and samples with high PKN2^KO^ scores (Fig. 3A, G). More detailed junctional analysis revealed that elevated PKN2^KO^ scores were specifically associated with gene programs favouring junction disassembly over assembly, with a preferential loss of tight junction over adherens junction components (Fig. 3G). Zona occludens-1 (ZO-1) immunostaining is commonly used to assess tight junction integrity in human intestinal tissue^23,24^. Immunostaining of colon sections revealed a marked reduction in ZO-1–positive area in both global and epithelial-specific PKN2^KO^ mice eight days after DSS treatment (Fig. 3I-K). Reduced ZO-1 staining was also evident in non-DSS treated mice two weeks following PKN2 deletion, although less pronounced at 15-days following DSS exposure (Supplementary Fig. S4H, I).

In line with impaired junctional stability, PKN2^KO^ organoids exhibited a strong enrichment of epithelial–mesenchymal transition programs, consistent with a shift toward wound-healing–like, junction-deficient epithelial states (Fig. 3A, G). PKN2-low colon (COAD) and rectal (READ) adenocarcinoma in TCGA data similarly showed significant enrichment in apical junction gene signatures, indicating this epithelial vulnerability may be preserved in epithelial-derived bowel cancers (Supplementary Fig. S4J, K). Together, these analyses demonstrate that loss of PKN2 recapitulates key features of human colitis and implicates PKN2 as a central regulator of epithelial junctional integrity that shapes disease severity and colitis-associated cancer risk.

### PKN2 loss sensitises intestinal epithelial tight junctions to inflammatory damage

Loss of junctional ZO-1 staining in vivo led us to investigate if PKN2 has a direct role in maintaining junction stability. PKN2 has previously been shown to associate with and regulate apical junction maturation in bronchial epithelial cells^25^. More recently, a study in MDCK-II cells also localised PKN2 to apical junctions as part of the Crumbs complex^26^. Staining of organoids confirms PKN2 is localised to ZO-1 positive apical tight junctions in primary colonic mouse epithelial cells (Fig. 4A). PKN2 was also localised with ZO-1 in human DLD1 and Caco2 colorectal cancer epithelial cell lines (Supplementary Fig. S5A, B). Knockdown of PKN2 using siRNA, or knockout with CRISPR-Cas9, disrupted ZO-1-positive tight junction formation in DLD1 cells (Fig. 4B-E, S5C-F) while leaving E-cadherin positive adherens junctions largely intact (Fig. 4F), indicating a selective requirement for PKN2 in tight junction formation.

**Fig. 4.**
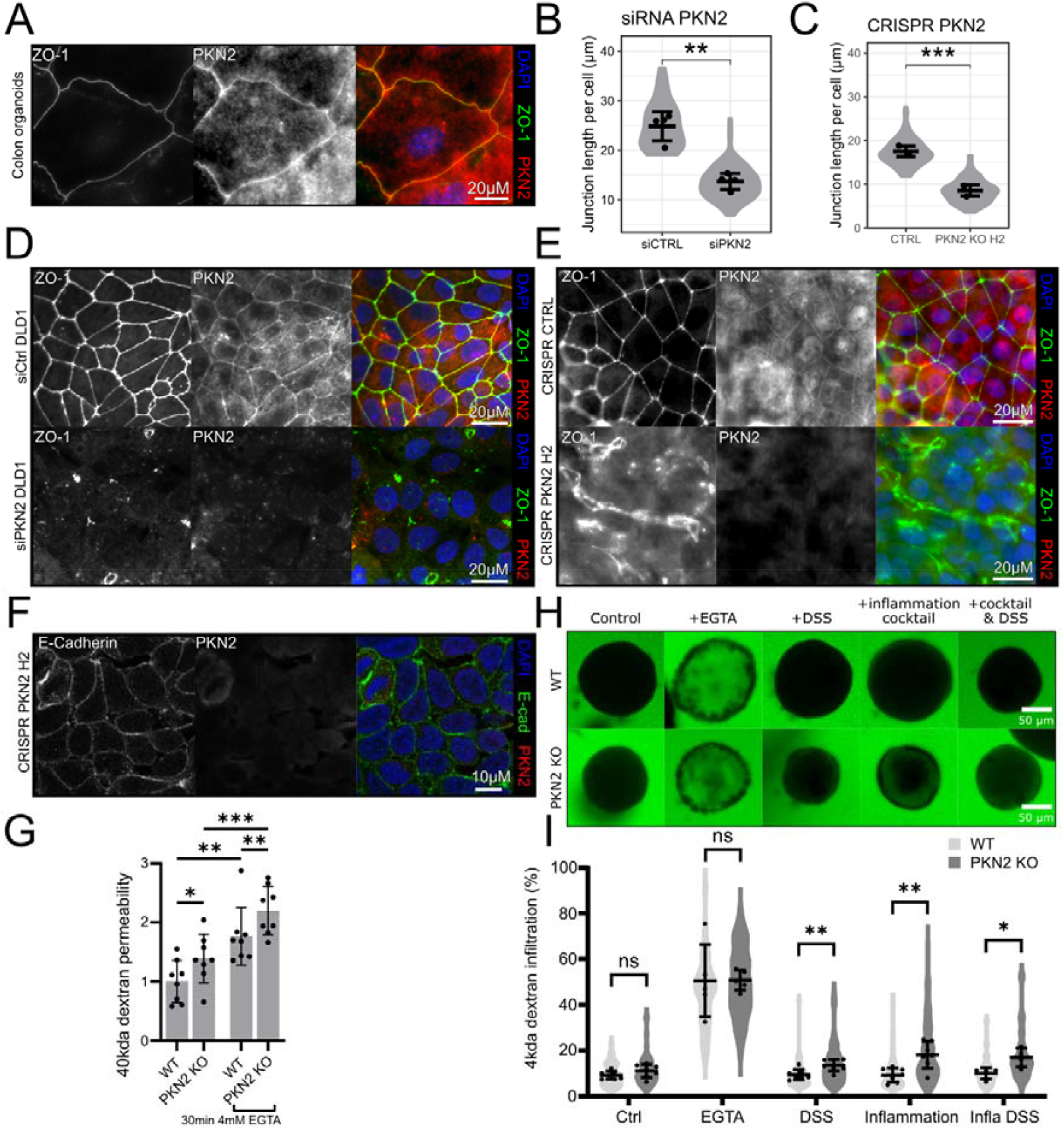
PKN2 loss sensitises intestinal epithelial tight junctions to inflammatory stimulation. **(A)** Immunostaining PKN2 and ZO-1 in C57BL6 colonic organoids cultured in 2D. **(B-E)** Disruption of ZO-1 positive tight junction formation following PKN2 depletion by siRNA (B, D) or CRISPR (C, E). Tight junction integrity was quantified as average length of junctional ZO-1 per cell (n=4, unpaired, two tailed t-test). **(F)** E-cadherin staining of adherens junctions in CRISPR PKN2 knockout DLD1 cells. **(G)** Permeability of CRISPR PKN2^KO^ DLD1 monolayers to 40kda FITC-dextran on 0.4µM Transwells treated with vehicle or 30mins 4mM EGTA (n=8; Fisher’s LSD test). **(H, I)** Permeability of 3D PKN2^KO^ organoids to 4kda FITC-dextran following treatment with 48h 10μg/ml DSS and/or 96h inflammatory cocktail (LPS, IL-1β, IL-6 and TNF-α)(n=7; Tukey’s test).

Functionally, tight junctions in colonic epithelia are essential for preventing the influx of luminal pathogens and toxins into the mucosa^27^. The inflammatory phenotypes observed in PKN2-deficient mice are therefore consistent with a sensitisation of epithelial tight junctions following DSS induced injury. In support of this, Functional Transwell™ assays demonstrate PKN2 knockout increases DLD1 epithelial monolayer barrier permeability to fluorescent 40kda FITC-dextran without compromising cell viability (Fig. 4G; S5G). In 3D organoid models, loss of PKN2 similarly resulted in an increase in permeability to fluorescent 4kda dextran, but only upon the administration of DSS or an inflammatory cocktail (Fig. 4H, I). Together, these results demonstrate that epithelial PKN2 is dispensable for homeostatic junction maintenance but is essential for preserving tight junction resilience under inflammatory stress, mirroring our in vivo observations.

### PKN2 loss disrupts linear actin regulation during tight junction assembly

To define how PKN2 contributes to tight junction assembly, we focused on the DLD1 epithelial model in which PKN2 function is sensitised and non-redundant. Imaging of junction formation in sparse cultures revealed that PKN2 is recruited to nascent cell-cell contacts together with F-actin and ZO1 (Fig. 5A). To test whether PKN2 kinase activity is required for junction formation we generated tetracycline-inducible DLD1 cell lines expressing either wild-type (iDLD1 WT-PKN2) or kinase-dead PKN2 (iDLD1 KD-PKN2 D782A). Induction of KD-PKN2 in iDLD1 cells exerted a dominant negative effect, impairing ZO-1 positive tight junction assembly (Fig. 5B, C, S5H, I). Conversely, induction of WT-PKN2 in iDLD1 cells had minimal impact on ZO-1-positive tight junction formation (Fig. 5D, S5J-L). These findings demonstrate that PKN2 catalytic activity is essential for its role in tight junction biogenesis. To explore potential substrates, we analysed phospho⍰proteomics datasets from PKN2⍰perturbed HEK293 cells ^28^. Perturbation of PKN2 altered phosphorylation states across a broad set of junctional and actin⍰associated proteins, including ZO⍰1 and PARD3 (Fig.⍰5E). This suggests that PKN2 orchestrates tight junction formation through a multi⍰layered phosphorylation programme targeting both tight junction regulators and the actin cytoskeleton.

**Fig. 5.**
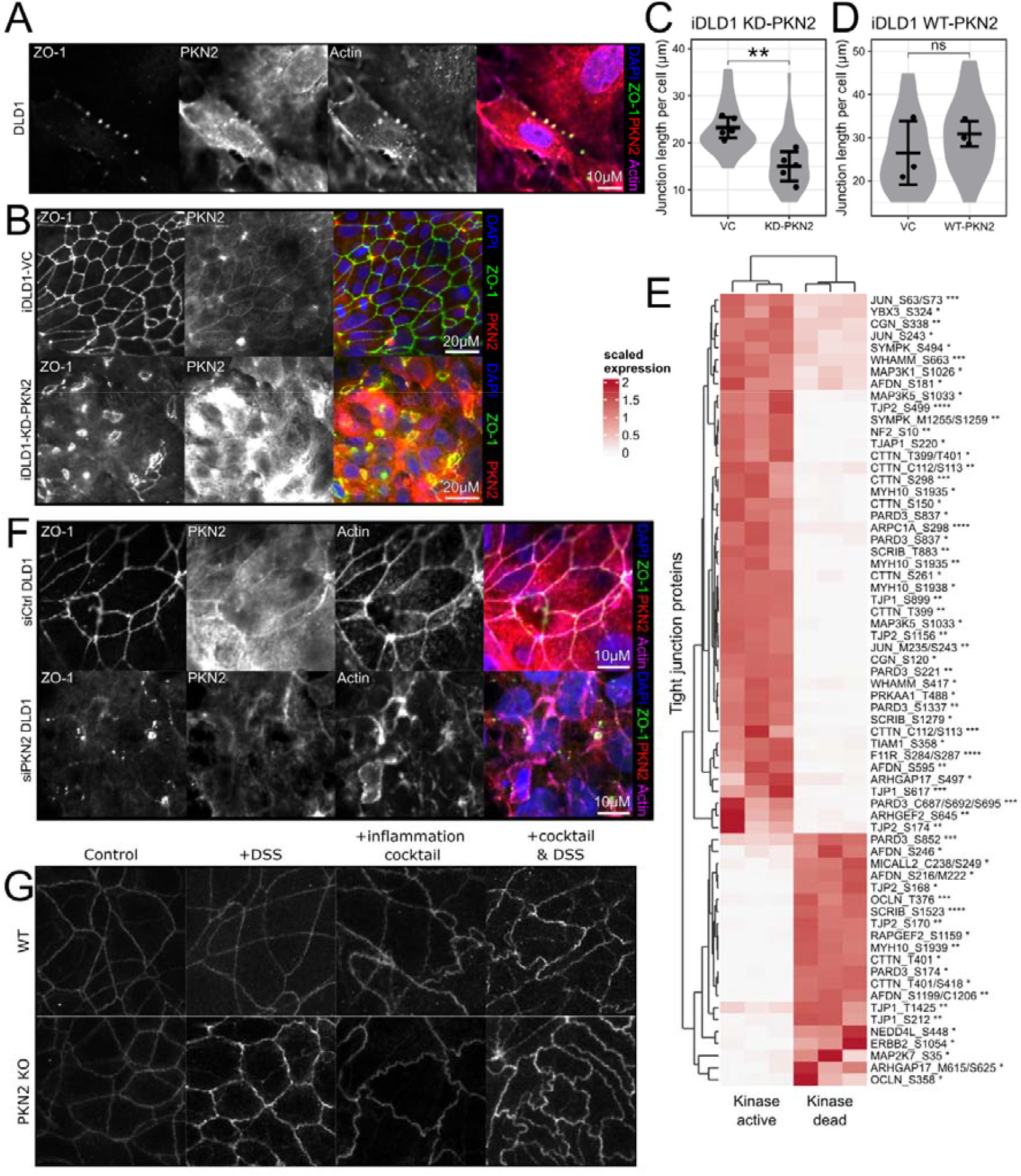
PKN2 loss sensitises intestinal epithelial tight junctions to inflammatory stimulation. **(A)** PKN2 localises to early nascent cell-cell contacts in sparse cultures of DLD1 cells. **(B-D)** ZO-1 positive tight junction formation following overexpression of a kinase-dead PKN2 (B, C) or overexpression of wild-type PKN2 (D). Tight junction integrity was quantified as average length of junctional ZO-1 per cell (C: n=4, D: n=3, unpaired, two tailed t-test). **(E)** Heatmap of significant changes to the phosphoproteome of junctional proteins with overexpression of kinase dead versus a kinase active PKN2 in HEK293 cells. **(F)** Disruption of peri-junctional actin belt formation following depletion of PKN2 via siRNA in DLD1 cells. **(G)** Changes to tight junction tortuosity of 3D PKN2^KO^ organoids to 4kda FITC-dextran following treatment with 48h 10μg/ml DSS and/or 96h inflammatory cocktail (LPS, IL-1β, IL-6 and TNF-α).

PKN2 has previously been implicated in the formation of linear actin bundles, with PKN2 depletion causing a failure in stress fibre assembly, structures that share architectural features with the circumferential junctional actin belt ^29^. Staining of F-actin using Phalloidin in PKN2 depleted DLD1 cells shows that loss of PKN2 disrupts the formation of the linear actin ring where actin structures appear miss-localised and disorganised (Fig. 5F). Similarly in 3D organoid cultures, PKN2 knockout produced a ruffled, tortuous junction phenotype, particularly following inflammatory stimulation (Fig. 5G). Increased junctional tortuosity is characteristic of impaired actomyosin tension along the apical actin belt, further supporting a role for PKN2 in organising the contractile linear actin architecture required for tight junction maturation. ZO-1 staining in PKN2 KO organoids, particularly following DSS treatment, is also less linearly distributed along the junctions consistent with an impairment of appropriate ZO-1 condensation. Together, these findings position PKN2 as a central organiser of the peri⍰junctional actin architecture that is essential for tight junction resilience under stress.

## Discussion

Our work identifies *PKN2* as a critical regulator of tight junction formation in the colon, protecting the bowel against inflammatory damage and colorectal cancer initiation. *PKN2*-targeted mice and primary organoids largely maintain morphologically intact tight junctions under homeostatic conditions, but significant defects emerge following epithelial injury, suggesting functional redundancy for normal tissue homeostasis. This concurs with knockout models of other core tight junction components, including O*CLN, JAM-A, PRKCI*, and *ZO-1*, which all maintain homeostatic junctions at baseline but exhibit heightened sensitivity to inflammation and colitis ^23,30-34^. These findings support the notion that homeostatic tight junctions are remarkably resilient to the loss of individual components, yet each contributes critically to maintaining or re-establishing barrier integrity following stress or injury.

Mechanistically, we identify PKN2 as a regulator of the peri⍰junctional actin belt, the linear actomyosin structure that provides the mechanical template for ZO⍰1 condensation and tight junction elongation ^35^. Since ZO⍰1 binds and polymerises along these linear actin tracks, appropriate belt formation is essential for anchoring tight junctions to the cytoskeleton and directing ZO-1 condensation along cell-cell boundaries. The linear actin bundles that form the circumferential junctional actin belt share core architectural features with stress fibres; both consist of parallel, unbranched but crosslinked F⍰actin filaments that incorporate non⍰muscle myosin II to generate contractile tension ^29^. Consistent with this shared architecture, PKN2 depletion or overexpression of a kinase dead PKN2 leads to the loss of stress fibres in multiple mesenchymal cell models, even under TGF-β stimulation ^19,36^.

PKN2 was previously found in proximity to Pals1 and co-immunoprecipitates with PatJ, both components of the Crumbs complex that reside immediately apical to mature tight junctions ^26^. Notably, depletion of PALS1 or PATJ similarly disrupts ZO⍰1 elongation and actin belt organisation, a phenotype attributed to impaired recruitment of the RhoA activator p114RhoGEF at nascent junctions^37,38^. Given that PKN2 is a RhoA⍰regulated kinase, our findings support a model in which PKN2 acts downstream of Crumbs⍰complex–mediated RhoA activation to assemble the linear actin bundles of the peri⍰junctional belt ^25^. In this framework, PKN2 promotes the transition from nascent cell–cell contacts to mature tight junctions by enabling the actin⍰dependent polymerisation and stabilisation of ZO⍰1.

The requirement for the kinase activity of PKN2 in junction formation suggests there are substrates for PKN2 in the junction and actin networks. Previous research identified PKN is a direct interactor of α-actinin and the loss of PKN2 reduces Palladin phosphorylation, implicating PKN2 in the regulation of two principle crosslinkers of linear actin bundles ^18,39^. In parallel, PKN2 is known to phosphorylate and inhibit cortactin, and potentially filamin A, thereby suppressing Arp2/3⍰dependent branched actin networks and reducing competition for formin⍰mediated linear actin polymerisation ^18,36^. PKN can also directly regulate actomyosin tension where it phosphorylates and increases the inhibitory effects of CPI-17 on myosin phosphatase activity ^40^. There may also be a broader phospho⍰regulatory role consistent with the widespread changes observed in ZO⍰1, Par3, and other junctional proteins upon PKN2 perturbation. Together, these findings support a model in which PKN2 coordinates multiple cytoskeletal and junctional pathways to drive the assembly, elongation, and stabilisation of the peri⍰junctional actin belt during tight junction maturation. ^26^

Intriguingly, the H.pylori bacteriotoxin CagA, which is known to localise to and disrupt junction formation and cell polarity^41,42^, directly binds to and inhibits the kinase activity of PKN2^43,44^. Our research suggests that inhibition of PKN2 by CagA, may directly contribute to H.pylori pathogenesis and promotion of colorectal and gastric cancer. Previous research also identified that PKN1 and PKN2 negatively regulate pyrin inflammasome activation in epithelial cells, and their combined knockdown leads to spontaneous IL-1β release, a cytokine known to increase epithelial permeability in colorectal cancer cell lines^45,46^. However, despite the pro-inflammatory role of IL-1β, multiple knockout models lacking inflammasome components show worsened colitis and AOM-DSS induced tumour burden^47-50^. Furthermore, knockout of PKN2 alone in organoid cultures did not significantly alter IL1B expression, suggesting that the tumour-suppressive effects of PKN2 are not primarily mediated through inflammasome dysregulation but instead through direct effects on tight junctions.

PKN2 was identified as one of 25 high-confidence candidate genes within CCs3, a locus associated with the heightened sensitivity of A/J mouse strain to AOM/DSS tumorigenesis, compared to C57BL/6J^17^. Further, PKN2 is among 340 genes within the *Cdcs1* locus, which contributes to enhanced susceptibility to disease in multiple genetic colitis models^16,51-53^. In addition, PKN2 lies within 207 genes on the Scc7 locus, which confers susceptibility to DMH-induced sporadic colorectal cancer in the recombinant congenic strain CcS-3 compared with the more resistant BALB/c background^54^. This suggests PKN2 is also a candidate risk factor on human 1p in sporadic colorectal cancer, consistent with the poor prognosis observed in patients with 1p allelic loss. Here we provide definitive proof that loss of PKN2 alone is sufficient to confer sensitivity to both colitis and colitis-associated-colorectal cancer, establishing PKN2 as a key functional candidate on human chromosome 1p for susceptibility to IBD and colitis-associated-colorectal cancer. Beyond colitis, GWAS studies have linked PKN2 to an increased risk of irritable bowel syndrome (IBS) (rs10923043) ^55^, and PKN2 is among susceptibility loci for periodontitis (1 of 16, rs12032672) and rosacea (1 of 98, rs191805855); conditions similarly linked to tight junction instability in the gums and skin respectively ^56-58^.

Primary organoids from patients with IBD had a remarkable similarity in their expression profile to primary murine PKN2^KO^ organoids, and PKN2 was identified as one of 18 significantly downregulated genes in endoscopic biopsies from patients with IBD when compared to healthy controls, alongside other junctional components such as PRKCI and CLDN8 ^59^. This suggests deregulation to core junction components, including PKN2, is a key mechanism underlying human IBD pathogenesis. It also highlights PKN2 as a potential therapeutic target, where stimulating its activity could reinforce barrier integrity and limit disease progression, a hypothesis that warrants testing in preclinical models. Our PKN2 knockout models also provide a platform for discovering new genetic and pharmacological approaches to counter junctional destabilisation and improve outcomes in human colitis and related epithelial disorders.

## Methods

### PKN2 knockout mouse models

All animal procedures were approved by the Queen Mary University of London ethics committee and performed under the UK Home Office Animal and Scientific Procedures Act (1986). Both sexes were included, with no sex⍰dependent differences observed.

Conditional Pkn2^fl/fl^ mice were crossed with B6.129-Gt(ROSA)26Sortm1(cre/ERT2)Tyj/J (JAX:008463) or B6.Cg-Tg(Vil1-cre/ERT2)23Syr/J (JAX:020282) mice. Pkn2^fl/fl^ mice were generated as previously described^18^. PKN2 deletion was induced using five daily intraperitoneal injections of tamoxifen (100⍰mg/kg in corn oil), followed by a 20⍰day recovery period.

### AOM/1%DSS mouse models

Following PKN2 knockout, mice received 1x intraperitoneal injection of azoxymethane (10mg/kg in PBS, 0.45μm-filtered, Sigma Aldrich), followed 3 days later by 3x cycles of dextran sodium sulphate (1/2% DSS in drinking water; 5 days, refreshed day 2, 16day recovery; Fig. 1A; MP Biomedicals). Tumours were analysed 40 days after the final DSS cycle.

Disease activity was assessed using a composite score of weight loss, stool consistency, haematochezia and general condition (1-3 per parameter, max 12) ^21^. Mice were sacrificed immediately upon a score of 3 in either weight loss or general condition, or after 24h of unresolved score 3 for stool consistency or haematochezia. Scoring was performed semi-blind.

### DSS only mouse models

Mice received 5 days of DSS (1/2% in drinking water, refreshed day 2) and were sacrificed 3/10 days later.

### Histology and immunohistochemistry

Colons were excised, Swiss⍰rolled^60^, fixed, paraffin⍰embedded and sectioned. Slides were dewaxed, rehydrated, subject to antigen retrieval (sodium citrate pH 6.0 10mM) and blocked with 3% hydrogen peroxide (Thermo-Fisher). Slides were then stained with CD4 (Thermo-Fisher, 14-9766-82), CD8 (Thermo-Fisher, 14-0808-82), F480 (Abcam, MCA497R), Ki67 (GeneTex, GTX16667), CD45 (Biolegend, 103101) and ZO-1 (Thermo-Fisher, 617300) overnight at 4°C and detected using Universal Vectastain ABC kit with Dab substrate (Dako, K3468). Slides were counterstained using haematoxylin (Sigma Aldrich) and mounted in DPX (Sigma Aldrich). Alcian Blue staining used 8GX 1% in 3% Acetic Acid (RRSK400-100).

Slides were scanned using a Nanozoomer-S210 (Hamamatsu) and analysed using QuPath (v0.5.1) ^61^. Erosion and submucosa length were averaged per mouse across multiple sections. Immune infiltrates were quantified through positive cell detection, alcian blue staining by random forest–based area quantification, and ZO-1 depletion by DAB intensity thresholding within an 18µm luminal region.

### PKN2 knockout organoid production and culture

Crypts of 12-week-old wild-type C57BL/6 and *Pkn2*^*fl/fl*^*;Rosa26-CreER*^*T2*^ mice were isolated by 2mM EDTA dissociation and embedded in growth-factor-reduced Matrigel (Corning) as described^62^. Organoids were cultured in Advanced DMEM/F-12 (Gibco), plus 2mM L-glutamine (Gibco), 1mM N-acetyl-l-cysteine (Sigma), 10mM HEPES (Sigma), B-27 (Gibco, 50X), N-2 (Gibco, 100X), HyClone Penicillin/Streptomycin (Cytiva), 10mM nicotinamide (Merck), 50ng mL−1 mEGF (PeproTech) and 25% WRN-conditioned media at standard conditions (37°C, 5% CO_2_). PKN2 knockout was induced by 4-hydroxytamoxifen (2µM TAM, 7-day treatment; Merck) and passaged every 7–10 days by mechanical dissociation. WRN-conditioned medium was generated from L-WRN cells as previously described^63^.

### Cell lines

DLD1 (ATCC), HEK293T (ATCC), Caco-2 and L-WRN cells were maintained in DMEM plus 10% FBS and 100U/ml penicillin/streptomycin (Gibco) under standard conditions (37°C, 5% CO_2_). Cell lines were routinely passaged using trypsin–EDTA (Gibco) and cryopreserved in 10%FBS/DMSO. Caco-2 were gifted from Egle Solito and L-WRN cells^63^ from Chris Tape.

### CRISPR-Cas9 PKN2 knockout cell lines

Human PKN2 targeting (CTTTGACGTGGACTTAGTGT-Exon 6) and non-targeting (GTATTACTGATATTGGTGGG) gRNA were cloned into the Lenti-Cas9-gRNA-GFP plasmid (Addgene #124770; gifted by Jason Sheltzer) as described^64^. Lentiviral particles were generated in HEK293T cells using a calcium⍰phosphate–based transfection approach (ProFection). Viral supernatants were collected and clarified before addition to DLD1 cells for 24h (50% media mix with 8µg/mL polybrene). GFP-positive cells were isolated by flow cytometry (ARIA Fusion).

### Inducible DLD1 cell lines

Stable inducible DLD1 (iDLD1) cells were produced by the integration of wild-type or kinase-dead PKN2 constructs (pcDNA5/FRT/TO/PKN2(WT/D782A)) in DLD1-FRT cells containing the Flp-In™ T-REx™ system (Invitrogen) (gifted by Stephen Taylor). After antibiotic selection (hygromycin (100μg/ml; Invitrogen)/blasticidin (10μg/ml; Thermo-Fisher)) for 30 days, transgene expression was induced with tetracycline (500ng/ml; Sigma).

### PKN2 siRNA knockdown

DLD1 cells were transfected with 20nM siRNA control (siCtrl #1) or PKN2 (siPKN2 #9, 10 & 11) pools (GE HealthCare) using Lipofectamine™ LTX (Thermo-Fisher) according to manufacturer’s instructions.

### Immunofluorescence

Cells or organoids were seeded onto coverslips, Nunc Lab-Tek II 8 well Chamber Slides (Thermo-Fisher) or 96-well PhenoPlates (Revvity). Cells or organoids were fixed with 4% PFA (organoids 2h, cells 15min; Thermo-Fisher), permeabilised with 0.2% Triton X-100 (15min; Sigma), blocked with 5% BSA (1h; Sigma), and incubated with PKN2 (R&D, MAB5686), ZO-1 (Invitrogen, 61-7300) or E-cadherin (Cell Signaling Technology, 3195T) overnight at 4 °C. PKN2 signal was amplified using Alexa Fluor™ 488 or 555 Tyramide SuperBoost (Invitrogen) according to the manufacturer’s instructions. Samples were incubated with DAPI, Phalloidin (Invitrogen, A22283), anti-Rabbit IgG Alexa Fluor™ 488/555/633 (Invitrogen, A-21206, A-31572, A-21071) or anti-Mouse IgG Alexa Fluor™ 488/555 (Invitrogen, A-11001, A-31570) and mounted in MOWIOL or left in PBS, and imaged by LSM 880 confocal microscope (Zeiss) or high-content IN Cell Analyzer 6000 (Cytiva). Image analysis was performed using Fiji ImageJ (v2) or CellProfiler (v4.2.8).

### Cell junctional staining analysis

A CellProfiler (v4.2.8) pipeline was assembled to perform junctional analysis. ZO1-stained junctions were enhanced using morph “openlines” (length 30) and “neurite-like” features enhanced with “tubeness” (smoothing 2). Enhanced junctions were smoothened (circular average filter & diameter of 4) and converted into junction objects using global robust background thresholding. Objects were filtered by minimum major axis length of 5.4µm to remove punctate artefacts and measurements taken. ZO1 junctional intensity was measured by masking the original image with the junction objects.

### Epithelial barrier permeability assays

DLD1 cells were seeded onto 0.4µm Pore Transwells (Corning) and cultured for 7 days under standard conditions. Monolayers were treated with control medium, 2h 50mg/ml DSS or 30mins 4mM EGTA, washed, and permeability assessed using 40kda FITC Dextran (5mg/ml in Phenol red-free DMEM, Merck). Flow-through samples were collected after 3h and measured at 485/535nm on FLUOstar OPTIMA microplate reader (BMG LABTECH).

Organoids embedded in Phenol Red-Free Matrigel (Corning) on a 96-well PhenoPlate (Revvity) were treated with either 2h 4mM EGTA, 96h inflammatory cocktail ((100ng/ul LPS (Sigma), 30ng/ml of IL-1β, IL-6, TNF-α (Gibco)), 48h 10μg/ml DSS or a combination thereof. Permeability was assessed with 4kda FITC Dextran (Merck). After 2h, images were taken by LSM 880 confocal microscope (Zeiss) and internalised dextran (%) was calculated using Fiji (v2).

### Western blotting

Powdered snap frozen tissue or cultured cells were lysed in NuPAGE LDS (Invitrogen) with 50mM DTT (Sigma-Aldrich), sonicated, and boiled at 95°C for 5mins. Protein concentration measured by DC protein assay kit (Bio-Rad), separated by SDS–PAGE (4–12% Bis-Tris gels (NuPAGE)) and transferred to nitrocellulose (Amersham) or PVDF (Millipore) membranes overnight (4°C). Membranes were blocked in 3% BSA TBS–Tween (0.1%) (Sigma) and incubated overnight (4°C) with primary antibodies, PKN2 (R&D MAB5686), GAPDH (Santa-Cruz SC-25778) or HSC70 (Santa-Cruz SC-7298), followed by HRP-conjugated secondary antibodies (GE Healthcare NXA931) and chemiluminescent detection (Luminata Forte/ Crescendo (Merck)).

### PCR genotyping

DNA was extracted from EGTA-dissociated colon crypts (tamoxifen treated *Pkn2*^*fl/fl*^*;Vilin-CreER*^*T2*^), organoids (QIAamp mini-DNA kit (Qiagen)) and FFPE sections (GeneJET FFPE DNA Purification Kit (Thermo-Fisher)). Multiplex PCR was performed (DNA using Platinum™ II Hot-Start PCR Master Mix (Thermo-Fisher)) using primers spanning the PKN2 alleles (GCATGCTGAGGATGTCACCG, GAGGGAAAATGGGAGAGG and GCAGTGGTGAAGTCACATAC). Samples were run alongside a 50bp ladder (NEB), on a 6% TBE gel (Thermo-Fisher), incubated with GelGreen (Merck) and imaged on a ChemiDoc-MP (Bio-Rad).

### Standard RNA sequencing

RNA was extracted using the Qiagen RNAeasy mini kit (QIAGEN) and quality assessed using a 5300 Fragment Analyzer (Agilent Technologies). Poly(A) selection Library preparation and paired-end RNA sequencing was performed by GENEWIZ (Azenta) using NovaSeq™ 6000 (Illumina). Trimming, aligning to GRCm39 and counting was performed using trimmomatic (v0.36), STAR (v2.7.9), samtools (v1.19.2) and subread (v2.0.6) using annotation files from GENCODE (M34).

### Smart-3SEQ RNA sequencing

Laser capture microdissection was used to isolate epithelial regions from FFPE sections, followed by Smart-3SEQ library preparation as previously described^22,65^. RNA quality was assessed using a Tapestation Fragment Analyser (Agilent Technologies) and sequenced using the Illumina NextSeq 2000 at the Genome Facility (ICR). FASTQ were processed as above with the addition of deduplication using umi_tools (v2.0.6).

### RNA sequencing analysis

All analysis was performed using custom R (V4.4) or bash scripts available on GitHub (https://github.com/jackchenry/PKN2_in_CA-CRC). Differential expression analysis was performed using DeSeq2 (v1.44) and gene-set enrichment analysis using fgsea (v1.30) with gene-sets from the molecular signatures database (v2024.1.Hs; v2024.1.Mm). Results from differential gene expression analysis, gene set enrichment analysis, raw count matrixes and sample annotations are available in Data S1. Visualisations were produced with ggplot2 (v3.5.1), ComplexHeatmap (v2.20) and GraphPad Prism (v10.1.6).

### External data and integrative analysis

Public RNA datasets were obtained from the Genomic Data Commons data portal (https://portal.gdc.cancer.gov/) and Gene Expression Omnibus under accessions GSE102746 ^66^, GSE193677 ^67^, GSE206285 ^68^, GSE109142 ^69^, GSE117993 ^69^, GSE207022 ^70^, GSE16879 ^71^, GSE59071 ^72^, GSE95437 ^73^, GSE235236 ^74^, GSE243625 ^75^, GSE53306 ^76^, GSE52746 ^77^. Known cancer genes were obtained from COSMIC. The PKN2^KO^ signature was created using organoid differential expression and GSVA. Immune infiltrate deconvolution was performed as previously described ^78^. Single-cell RNA data was obtained from PREDICT ^79^. PKN2 proteomic data was obtained from the source publication ^28^. PrismEXP prediction results on ARCHS4 data was obtained for PKN2 from the ARCHS4 data portal (v2.5, https://archs4.org/)^80^

### Statistical analysis

All statistical analysis was performed in R (V4.4) or GraphPad Prism (v10.1.6). Different statistical tests are used throughout the study as indicated in figure legends. A minimum of n = 3 independent experiments was used, as indicated in the figure legends. P < 0.05 was statistically significant.

## Abbreviations

AOM: Azoxymethane
ATCC: American Type Culture Collection
BSA: Bovine serum albumin
CCs3: Colon cancer susceptibility locus on mouse chromosome 3
Cdcs1: Cytokine deficiency-induced colitis susceptibility-1
COAD: Colon adenocarcinoma (TCGA)
CreERT2: Tamoxifen-inducible Cre recombinase
DAPI: 4⍰,6-diamidino-2-phenylindole
DAI: Disease Activity Index
DMEM: Dulbecco’s Modified Eagle Medium
DMSO: Dimethyl sulfoxide
DSS: Dextran sodium sulphate
EDTA: Ethylenediaminetetraacetic acid
EGTA: Ethylene glycol-bis(β-aminoethyl ether)-N,N,N⍰,N⍰-tetraacetic acid
FBS: Fetal bovine serum
FFPE: Formalin-fixed paraffin-embedded
FITC: Fluorescein isothiocyanate
GEO: Gene Expression Omnibus
GSVA: Gene Set Variation Analysis
GSEA: Gene Set Enrichment Analysis
H&E: Haematoxylin and eosin
HET: Heterozygous
HRP: Horseradish peroxidase
IBD: Inflammatory bowel disease
IBS: Irritable bowel syndrome
IL-1β: Interleukin-1 beta
IL-6: Interleukin-6
iPKN2^KO^: Inducible PKN2 knockout
JAM-A: Junctional adhesion molecule A
KD: Kinase-dead
KEGG: Kyoto Encyclopedia of Genes and Genomes
KO: Knockout
LPS: Lipopolysaccharide
PBS: Phosphate-buffered saline
PCA: Principal Component Analysis
PFA: Paraformaldehyde
PKN2: Protein kinase N2
PVDF: Polyvinylidene fluoride
READ: Rectal adenocarcinoma (TCGA)
SDS–PAGE: Sodium dodecyl sulphate–polyacrylamide gel electrophoresis
siRNA: Small interfering RNA
TAM: Tamoxifen / 4-hydroxytamoxifen
TCGA: The Cancer Genome Atlas
TBS: Tris-buffered saline
TNF-α: Tumour necrosis factor alpha
UMI: Unique molecular identifier
Vil1-CreERT2: Villin promoter-driven inducible Cre line
WT: Wild-type
ZO-1: Zona occludens-1

## Data availability

Data supporting the findings in this study are publicly available in supporting documents or Gene Expression Omnibus (GEO) under accession number GSE313373 and GSE313375. All custom analysis code used to generate results and figures is available at [https://github.com/jackchenry/PKN2_in_CA-CRC].

## Author Contributions

J.C.H. and A.J.M.C. conceptualised and developed the study. J.C.H performed the majority of methodology and formal analysis. A.C. contributed cell based and organoid studies. J.C.H. and P.H. conducted RNA sequencing, tissue staining and analysis with help from A.P., K.S. and M.L. J.C.H, J.S., S.M. and A.J.M.C. contributed to in vivo experiments. I.R., I.Q. and A.J.M.C. conducted mouse model development. M.J. and S.M. assisted with pathology and analysis. T.G., N.Q.M. and S.M. supported project development and data collection. J.C.H. and A.J.M.C wrote the manuscript with comments and approval from all the authors.

## Acknowledgements

We thank the core services at the Barts Cancer Institute, including Biological Services, Animal Technician Services, Pathology, Preclinical Imaging, and Microscopy. We are also grateful to Queen Mary’s Apocrita HPC facility, supported by QMUL Research-IT, and to the Phenotypic Screening Facility at the Blizzard Institute. We acknowledge the Genomics Facility at the Institute of Cancer Research. We also thank Professor Peter Parker and Ian Rosewell for their help in developing PKN2-targeted mice. Finally, we are grateful to the laboratories of Shinta Sato, and Egle Solito for supplying L-WRN and Caco2 cells.

This study was supported by the Medical Research Council (MR/X018997/1) and Cancer Research UK Centre grants to Barts Cancer Institute (C355/A25137 and C355/A29277) and the City of London Centre (C7893/A26233). Additional support was provided by grants from Worldwide Cancer Research/Pancreatic Cancer Research Fund (18-0713), Barts Charity (G-002149) and the Medical College of St Bartholomew’s Hospital Trust (P.H. Clinical PhD Fellowship).

