## Supplementary figures for "PKN2 regulates cell-junctions to limit colitis"

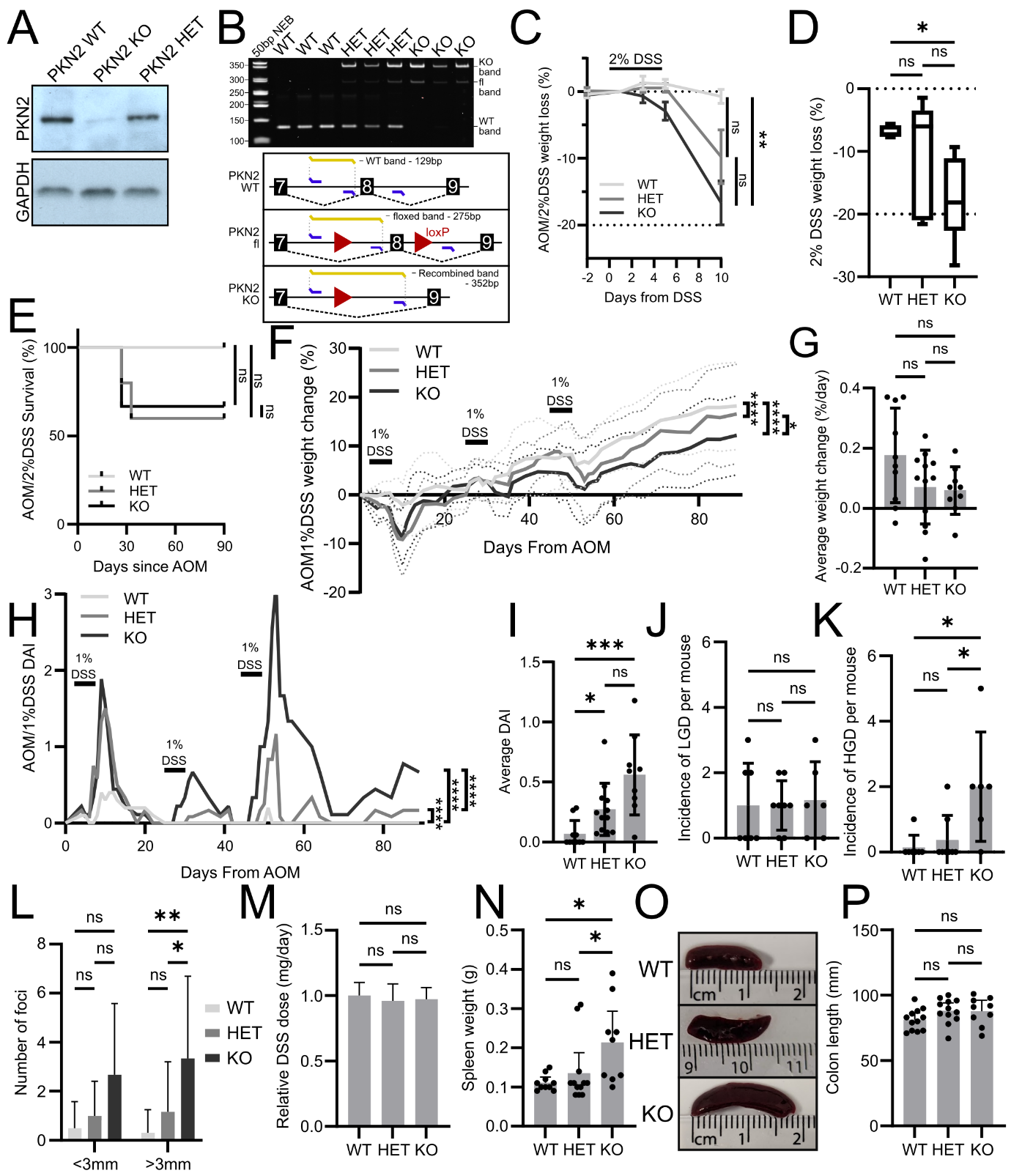

**Figure S1 - PKN2 loss sensitises mice to DSS treatment.** (**A**) PKN2 protein expression of intestinal tissue determined by western blot following the tamoxifen treatment regime. (**B**) PCR genotyping of intestinal tissue two weeks following the tamoxifen treatment regime of global-iPKN2^KO^ mice. Schematic describing the primer strategy and expected band sizes is shown in the bottom panel. (**C, D**) Average (C) and maximum (D) weight loss in response to the first 10 days of AOM/2%DSS treatment in WT, HET and PKN2 KO mice (WT: n=5, HET: n=5, KO: n=6, two-way repeated measures ANOVA (C) Dunn’s test (D)). **E)** Survival of WT, HET and PKN2 KO mice in response to 1 dose of 10mg/kg AOM followed by 5 days treatment with 100mg/kg tamoxifen and 1 round of 2%DSS for five days (WT: n=17, HET: n=18, KO: n=17; Log-rank test). (**F-I)** Average weight loss (F, G) and disease activity scores (H, I) across the second and third rounds of 1%DSS treatment in WT, HET and PKN2 KO mice (WT: n=10, HET: n=12, KO: n=9; two-way repeated measures ANOVA (F, H) or a Dunn’s test (G, I)). **(J, K)** Quantification of low (LGD, J) and high-grade dysplasia (HGD, K) in WT, HET and KO adenomas formed from the AOM/1%DSS treatment protocol (WT: n=7, HET: n=8, KO: n=6). (**L)** The average number of small (<3mm) and large (>3mm) diameter macroscopically visible adenomas from the AOM/1%DSS protocol across WT, HET and PKN2 KO mice (WT: n=10, HET: n=12, KO: n=9). (**M**) Relative average DSS dose measured through water bottle weight across WT, HET and PKN2 KO mice (WT: n=10, HET: n=12, KO: n=9). (**N-P**) Representative image of spleen size (O) and average weight (N) and average colon length (P) in WT, HET and PKN2 KO mice following the full the AOM/1%DSS protocol (WT: n=10, HET: n=12, KO: n=9; Tukey’s test).

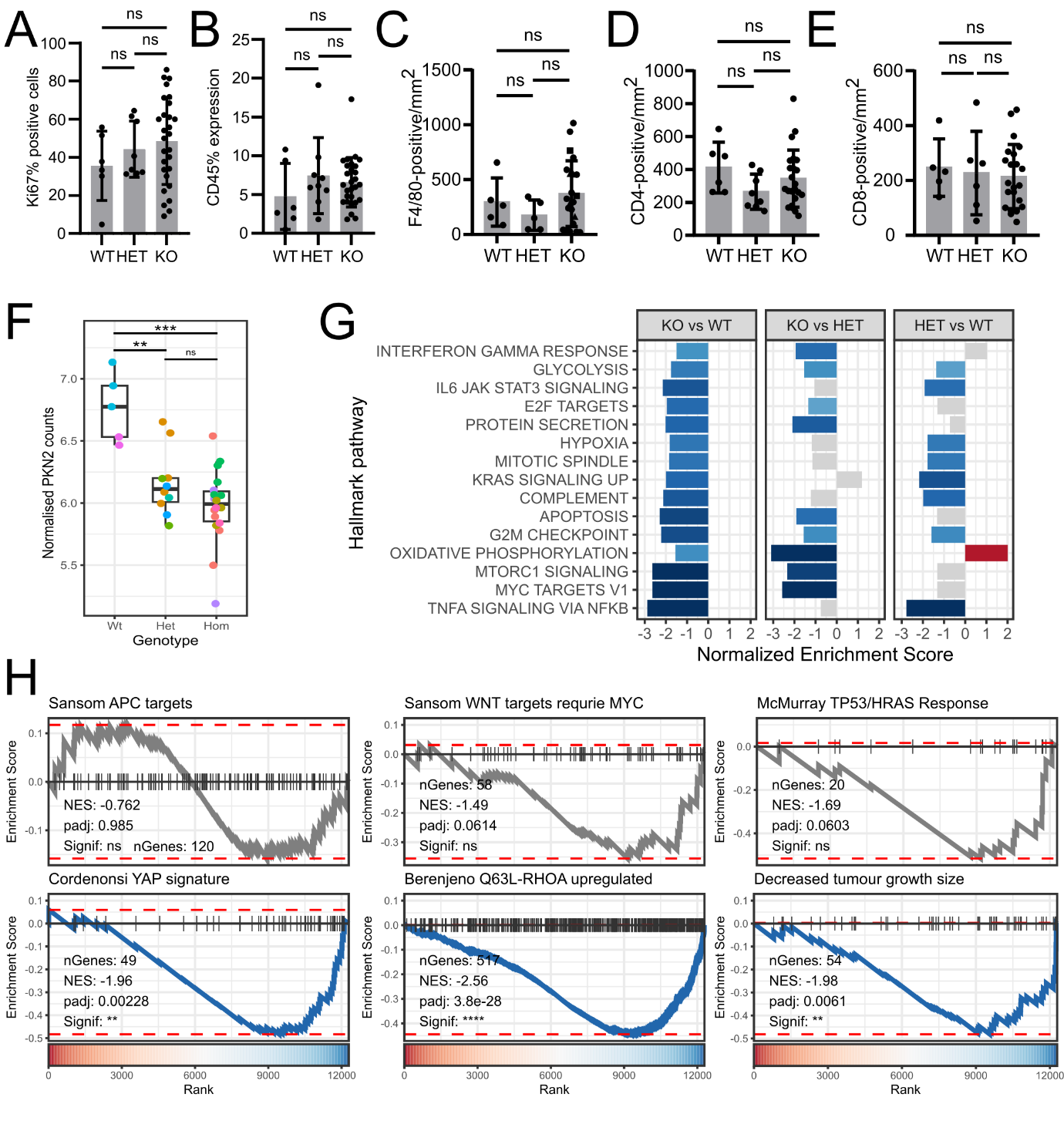

**Figure S2 - PKN2 status does not dictate tumour phenotype.** (**A-E**) Immunohistochemistry analysis of CD45 (A), F4/80 (B), CD4 (C) and CD8 (D) staining in WT, HET and PKN2 KO adenomas formed during the AOM/1%DSS protocol. (**F**) PKN2 expression from RNA sequencing analysis in adenomas by genotype. Statistical significance was determined using DESeq2 differential expression analysis (WT: n=5, HET: n=10, KO: n=18). Adenomas from the same individuals are coloured with the same colours. (**G**) Gene set enrichment analysis of KO vs WT, KO vs HET and HET vs WT adenomas for the most significant Hallmark gene sets. The top 15 most significant gene sets across all comparisons are shown. (**H**) Gene set enrichment analysis of KO vs WT for colorectal Sansom APC and WNT, McMurray TP53/HRAS, Cordenonsi YAP, Berenjeno Q63L-RhoA and mouse phenotypes decreased tumour growth size gene sets.

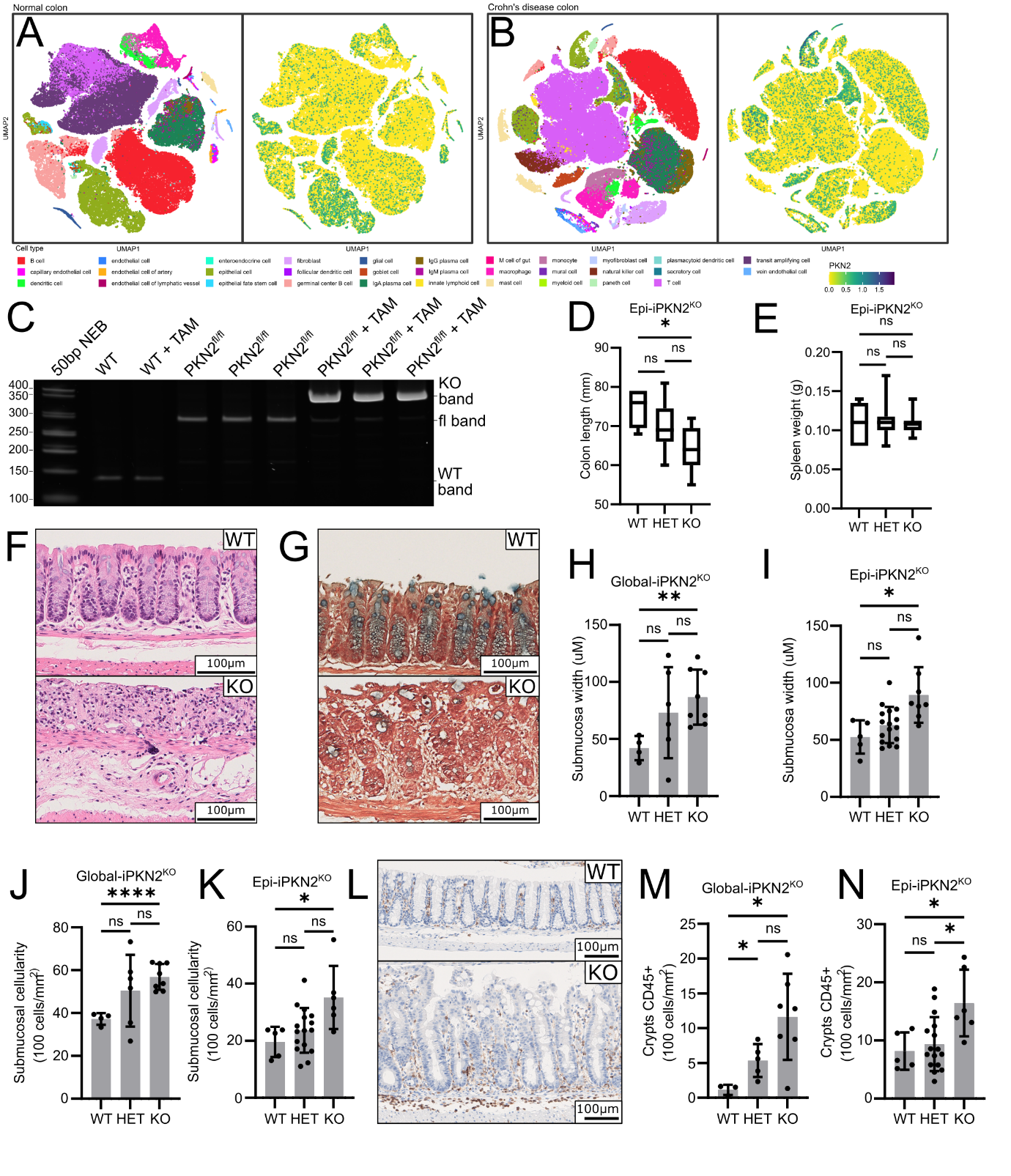

**Figure S3 - Epithelial specific PKN2 loss induces inflammatory damage.** (**A, B**) Single cell RNA sequencing analysis of samples from normal colon (A), Crohn’s disease patients (B). Cell subtypes are coloured in the left panel with PKN2 expression labelled in the right panel. (**C**) PCR genotyping of isolated colonic epithelial tissue two weeks following the tamoxifen treatment regime in Epi-iPKN2^KO^ mice. Diagram explaining band sizing is in Fig. S1C. (**D, E**) Colon length (D) and spleen weight (E) in Epi-iPKN2^KO^ mice following 5-days treatment with 2%DSS (WT: n=5, HET: n=16, KO: n=8; Dunnett’s test). (**F, G**) Epithelial erosion (F) and goblet cell depletion (G) indicated by alcian blue staining in Epi-iPKN2^KO^ mice following 5-days treatment with 2%DSS. (**H-K**) Average distal submucosa width (H, I) and cellularity (J, K) in global-iPKN2KO (H, J) and Epi-PKN2KO (I, K) mice following treatment with AOM/1%DSS (H, J) or 2%DSS (I, K). (H, J: WT: n= 4, HET: n=6, KO: n=8; Dunnett’s test), (I, K: WT: n=5, HET: n=16, KO: n=8; Dunnett’s test). (**L-N**) CD45+ immune cell infiltration in distal crypts and submucosa global-iPKN2KO (N) and Epi-PKN2KO (L, N) mice following treatment with AOM/1%DSS (M) or 2%DSS (L, N). (M: WT: n=3, HET: n=5, KO: n=7; Dunnett’s test) (N: WT: n=5, HET: n=16, KO: n=6; Dunnett’s test).

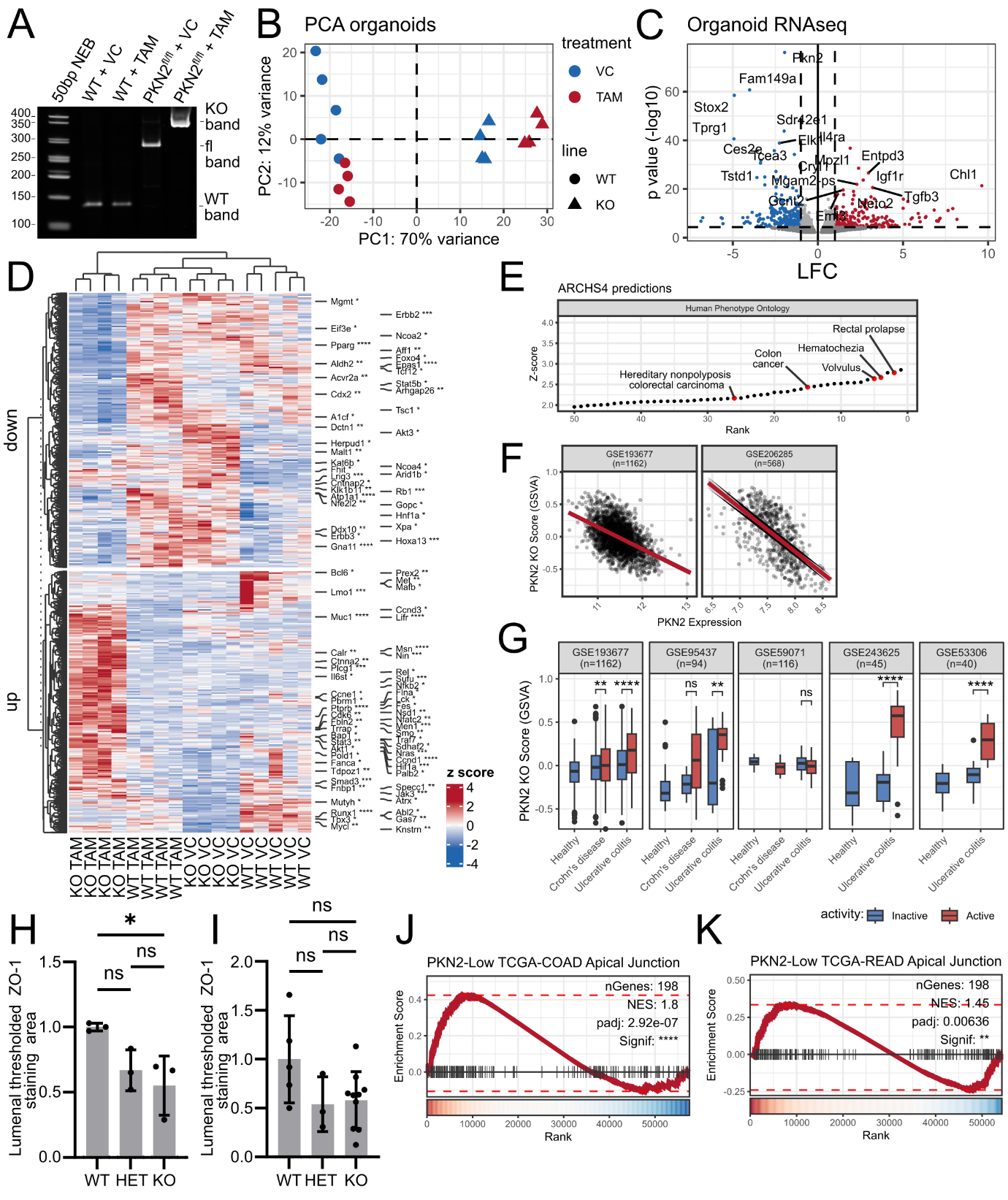

**Figure S4 - PKN2 loss alters gene expression, pathway enrichment, and junctional programs in cancer and colitis models.** (**A**) PCR genotyping of colonic organoids one week following the in vitro tamoxifen treatment regime. Diagram explaining band sizing is in Figure S1B. (**B**) Principal component analysis of RNA sequencing data from wild-type and Rosa^Cre/+^/PKN2^fl/fl^ primary mouse colon organoids treated with 4-OHT tamoxifen or a vehicle control. (**C**) Differential gene expression analysis identifying the most significantly differentially expressed genes between vehicle control and tamoxifen treated PKN2-KO organoids, accounting for the effects of tamoxifen treatment on wild-type organoids. (**D**) Significant differentially expressed genes between vehicle control and tamoxifen treated PKN2-KO organoids, accounting for the effects of tamoxifen treatment on wild-type organoids. Known cancer genes from the Cancer Gene Consensus are labelled. **E)** Top 50 ranked human phenotype ontology gene sets for predictions of PKN2 function by PrismEXP across the ARCHS4 compendium. (**F**) Scatter plots describing the relationship between PKN2 expression and PKN2 knockout score. (**G**) Differences in PKN2 KO score comparing patients with active and inactive disease. (**H, I**) Immunohistochemical analysis of distal colonic luminal epithelial ZO-1 staining in mice two weeks following PKN2 knockout induction (H) or 15 days following the initiation of 5 days of 1% DSS treatment (I) (H: n=9, I: n=18; Holm-Šídák's test). (**J, K**) GSEA of hallmark apical junction gene set in TCGA-COAD (J) and TCGA-READ (K) ranks created from comparing PKN2-low and PKN2-high patient primary tumours.

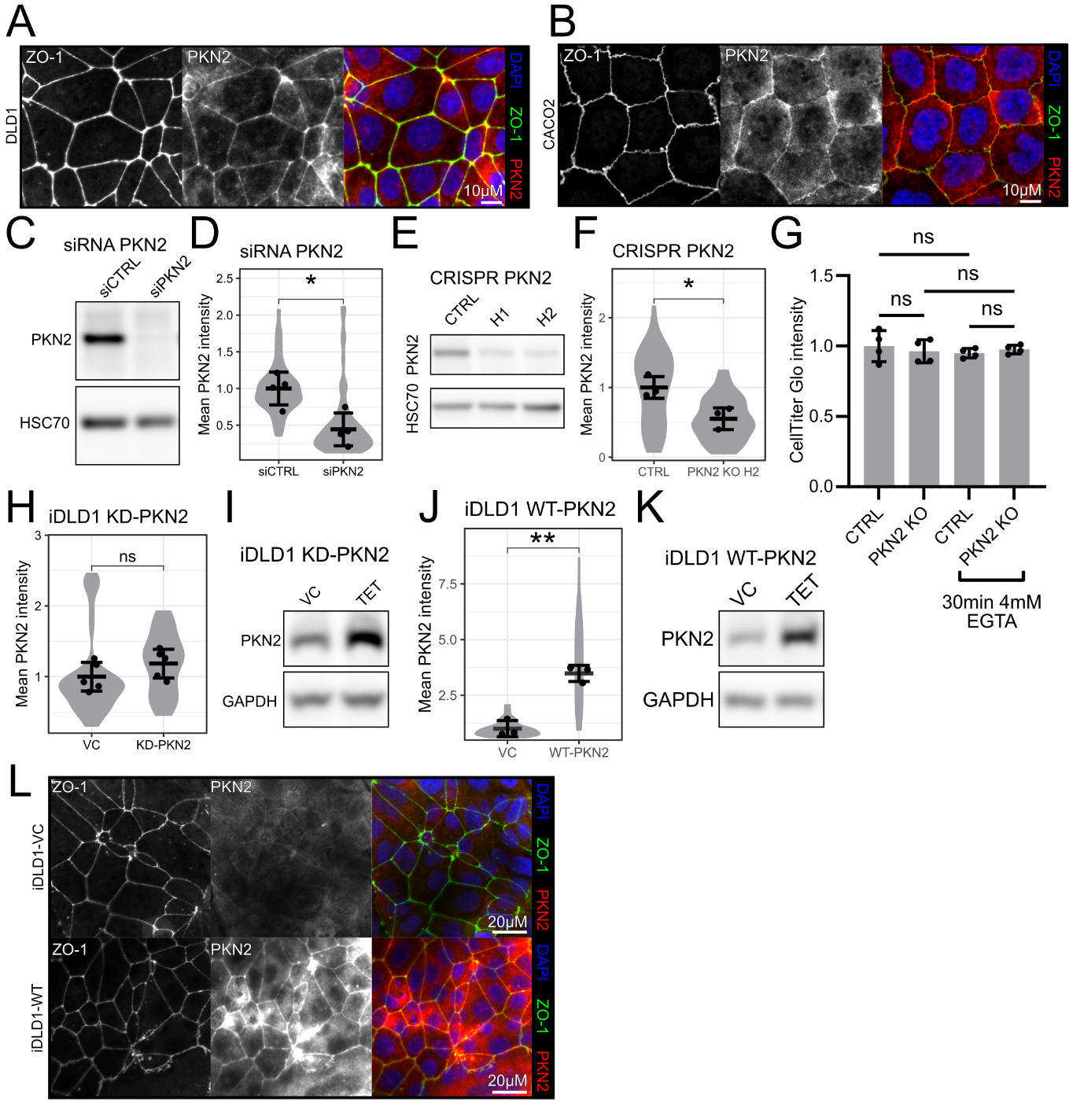

**Figure S5 - PKN2 regulates junctional ZO‑1 integrity in cells, organoids, and mouse colon epithelium.** (**A-B**) Colocalization staining of junctional PKN2 and ZO-1 in DLD1 (A) and CACO2 (B) cells. (**C-F**) Loss of PKN2 protein expression following PKN2 knockdown with a PKN2-targeting siRNA pool (C, D) or PKN2 knockout using CRISPR (E-F) by immunofluorescence and western blot analysis (n=4, unpaired, two-tailed t-test). (**G**) Difference in in CellTitre Glo intensity between CRISPR control and PKN2 KO gRNA DLD1 lines. (**H-K**) Gain of PKN2 expression following overexpression of a kinase-dead (KD, H, I) or wild-type (WT, J, K) PKN2 in the inducible DLD1 cell line (iDLD1) with and without 4mM EGTA treatment (H: n=5, J: n=3; unpaired, two-tailed t-test). (**L**) Junction morphology following overexpression of WT-PKN2 in the iDLD1 cell line.
